# Lack of Evidence that Ursodeoxycholic Acid’s Effects on the Gut Microbiome Influence Colorectal Adenoma Risk

**DOI:** 10.1101/237495

**Authors:** Talima Pearson, J. Gregory Caporaso, Monica Yellowhair, Nicholas A. Bokulich, Denise J. Roe, Betsy C. Wertheim, Mark Linhart, Jessica A. Martinez, Cherae Bilagody, Heidie Hornstra, David S. Alberts, Peter Lance, Patricia A. Thompson

**Author notes:** Authors share first authorship (Talima Pearson, J. Gregory Caporaso). Authors share senior authorship (Peter Lance, Patricia A. Thompson). Correspondence: J, Gregory Caporaso, PhD, Pathogen and Microbiome Institute, Flagstaff, AZ, 86011.; Peter Lance, MD, University of Arizona Cancer Center, 1515 N. Campbell Avenue, Tucson, AZ, 85724. Patricia A. Thompson, PhD, Stony Brook School of Medicine, Stony Brook, NY, 11794. Author contributions: Conception, funding, and design: TP, JGC, DSA, PL, PAT. Data generation: MY, CB, HH. Analysis/interpretation: TP, JGC, NAB, DJR, BCW, JAM, DSA, PL, PAT. Data access: Raw sequence data is currently being deposited in the Qiita microbiome meta-analysis repository, and the ENA sequence read archive. Accession numbers will be included in the final version of this manuscript, after they have been generated. For review purposes, data can be anonymously accessed through dropbox using the following link: https://www.dropbox.com/sh/49115v1mukwjtc3/AABdB8hx33ogeheILLmvom9na?dl=0.

## Abstract

**Objective:** We previously reported that Ursodeoxycholic acid (UDCA), a therapeutic bile acid, reduces risk for advanced colorectal adenoma in men but not women. Interactions between the gut microbiome and fecal bile acid composition as a factor in colon cancer neoplasia have been postulated but evidence is limited to small cohorts and animal studies.

**Design:** Using banked stool samples collected as part of a phase III randomized clinical trial of UDCA for the prevention of colorectal neoplasia, we compared change in the microbiome composition after 3 years intervention in a subset of participants randomized to 8–10 mg/kg of body weight UDCA (n=198) to placebo (n=203). UDCA effects on the microbiome, sex and adenoma outcome were investigated.

**Results:** Study participants randomized to UDCA experienced compositional changes in their microbiome that were statistically more similar to other individuals in the UDCA arm than to those in the placebo arm. This change reflected an UDCA-associated shift in microbial community distance metrics (P <0.001), independent of sex, with no evidence of UDCA effect on microbial richness (P > 0.05). These UDCA-associated shifts in microbial community distance metrics from baseline to end-of-study were not associated with risk of any or advanced adenoma (all P> 0.05) in men or women.

**Conclusion:** Despite a large sampling of randomized clinical trial participants, daily UDCA use only modestly influenced the relative abundance of microbial species in stool with no evidence for effects of UDCA on stool microbial community composition as a modifier of colorectal adenoma risk.

**SUMMARY:** *What is already known about this subject?:*

- Ursodeoxycholic acid (UDCA) is a therapeutic bile acid used in the treatment of primary biliary cirrhosis (PBC) and investigated for anti-cancer activity in the colon
- In humans, UDCA is produced in the colon from the conjugation of primary bile acids by intestinal bacteria
- Intestinal bacteria play a critical role in human intestinal health and disease including a hypothesized role in the development of colorectal cancer.
- UDCA was found to reduce the risk of more advanced colorectal adenoma with effects present in men but not women.
- Therapeutic UDCA was recently shown to reduce the extent of bacterial dysbiosis in patients with PBC

*What are the new findings?:*

- Among a population of patients with colorectal adenoma, low dose oral UDCA taken daily produced modest changes in fecal bacterial composition
- UDCA associated changes in the gut microbiome were similar in men and women.
- UDCA associated changes in the gut micobiome were not associated with risk of any or advanced colorectal adenoma in the patient population.

*How might it impact on clinical practice in the foreseeable future?:*

- These findings confirm effects of oral UDCA on the microbiome that may be beneficial for patients with PBC.
- These findings suggest that the anti-cancer effects of UDCA for colorectal adenoma prevention are not due to major effects of UDCA on the gut microbiome.

## INTRODUCTION

Western diet and lifestyle account for up to 80% of colorectal cancer (CRC) incidence.^1^ Numerous specific factors are proposed to explain this association, and their influence on the gut microbiome as a factor in CRC risk is a longstanding hypothesis.^2^ The interplay between gut bacterial composition and host epithelium is recognized in local immune function, metabolism, and host health, including an hypothesized role in susceptibility to gastrointestinal and other cancers.^3, 4^ Reported differences in the gut microbiome, including microbial community composition, between healthy and tumor tissues support disturbances in intestinal bacteria associated with CRC.^5^ This includes evidence of dense colonies of bacteria (i.e., biofilms) invading the mucus layer in association with colonic adenoma and cancers, particularly of the right colon, that *in vitro* exhibit tumor-promoting effects.^6^

Establishing a causal relationship between gut bacteria and colonic neoplasia has been elusive. The best evidence for an etiologic role for gut bacteria in CRC has been obtained from mouse model studies.^2^ For example, in the dextran sodium sulfate (DSS) inflammation-accelerated azoxymethane (AOM) mouse model of CRC, antibiotic treatment prior to and during AOM injection and throughout DSS treatment reduced tumor size and number.^7^ Further, stool and bedding from tumor-bearing mice transferred to germ-free mice treated with AOM/DSS increased tumor size and number. Interestingly, treatment with combination AOM/DSS also was shown to alter microbial community composition. Together, such findings support microbiome remodeling as an important component of tumor development and progression.

Several hypotheses are proposed to explain a role for gut bacteria in CRC, including the tumorigenic activity of secondary bile acids *[e.g*., deoxycholic acid (DCA)] produced by bacterial bile salt hydrolases in the large intestine.^2, 8–10^ Outstanding interest in a bile acid-CRC hypothesis led us to investigate ursodeoxycholic acid (UDCA): a therapeutic bile acid based on evidence of preventive activity in mouse models of colon carcinogenesis,^11^ favorable effects of UDCA on bile acid pools including DCA-lowering activity,^12^ reports of lower CRC risk in patients receiving UDCA for other indications^13, 14^ and recent evidence that dysbiosis in the gut microbiome of patients with primary biliary cirrhosis (PBC) may be modified by treatment with UDCA.^15^ In our phase III placebo-controlled, randomized trial of UDCA, we observed no effect of UDCA on adenoma overall at follow-up, but we noted reduction in adenoma with high-grade dysplasia.^16^ Subsequently, we showed reduced risk for large and advanced adenoma in men randomized to UDCA, and evidence for increased risk among younger and obese women, implicating sex as an important variable in UDCA activity.^17^ Since completing this trial, evidence for sexual dimorphism in bile acid metabolism in mice^18^ and bile acid effects on gut bacterial composition^19^ have emerged, prompting us to evaluate UDCA for its effects on the microbiome and adenoma outcomes, with consideration for sex. We used archival paired stool specimens from a subset of participants in the UDCA trial to test the effect of UDCA on the microbiome and conduct exploratory analyses to relate microbiome measures to adenoma outcomes.

## MATERIALS AND METHODS

### Study group, sample collection, study design

As part of the Arizona phase III placebo-controlled trial of 8–10 mg/kg of body weight UDCA for the prevention of colorectal adenomas, stool samples were obtained from participants who consented to fecal bile acid analysis.^20, 21^ Briefly, eligible individuals had at least one colorectal adenoma with a diameter of ≥3 mm removed during a colonoscopy within six months before registration. A total of 1,285 participants were randomized to UDCA (n = 661) or placebo (n = 624), of whom 1,192 (613 UDCA and 579 placebo) completed the trial. The primary trial endpoint was colorectal adenoma, defined as the occurrence of one or more adenoma or adenocarcinoma at colonoscopy performed ≥ 6 months after the qualifying colonoscopy. Advanced adenomas were defined as previously described as those with adenocarcinoma, high-grade dysplasia, villous/tubulovillous histology, or a diameter ≥ 1 cm.^17^ All stools passed over a 72-hour period were collected in a single metal container on ice. Pooled 72-hour samples were transported at 4°C to the laboratory where fecal solid was separated from fecal water as previously described.^20, 21^ Separated fecal water and solid stool were stored at –80°C for an average of 15 years until processing for microbial DNA.

For the current study, only participants with paired baseline (pre-intervention with UDCA or placebo) and end-of-study microbiome sequence data and adenoma outcome data were included. A total of 401 participants (198 UDCA and 203 placebo) with paired samples generated 802 total samples for analysis.

### DNA Extraction

DNA was extracted from thawed stool samples using the QIAamp DNA Stool Mini Kit protocol (Qiagen Inc., Valencia, CA) according to the manufacturer's instructions without modifications. Briefly, 200 mg of feces was placed in a sterile, round-bottom 2 mL tube containing 1.4 mL ASL lysis buffer. The homogenate was pelleted and incubated with InhibitEX to adsorb inhibitors. Proteinase K and Buffer AL were added to the supernatant to digest proteins. The DNA was bound to a spin column filter, and impurities were washed from the sample using 96–100% ethanol and proprietary Buffer AW2. All samples were eluted in 200 μL AE buffer and stored at –80°C until use in PCR.

### PCR and Sequencing

PCR of the V4 region of the 16S rRNA gene and sequencing were performed on the Illumina MiSeq platform following the original Earth Microbiome Project protocols (http://www.earthmicrobiome.org/protocols-and-standards/) originally described by Caporaso et al.^22^

### Bioinformatics

Microbiome bioinformatics were performed with QIIME^23^ 2 (https://qiime2.org/) 2017.4, a plugin-based system that, in some cases, wraps other microbiome analysis methods. Briefly, raw sequence data were demultiplexed and quality filtered using the q2-demux plugin followed by denoising with DADA2^24^ (via q2-dada2) to identify all observed amplicon sequence variants (ASVs)^25^ [i.e., 100% operational taxonomic units (OTUs)]. All ASVs were aligned with mafft^26^ (via q2-alignment) and used to construct a phylogeny with fasttree2^27^ (via q2-phylogeny). Alpha-diversity metrics (observed OTUs and Faith’s Phylogenetic Diversity^28^ – measures of microbiome richness) and beta-diversity metrics (weighted UniFrac^29^, unweighted UniFrac^30^, Jaccard distance, and Bray-Curtis dissimilarity – measures of microbiome composition dissimilarity) and principal coordinate analysis (PCoA) were estimated using q2-diversity after samples were rarefied (i.e., subsampled without replacement) to 900 sequences per sample. Taxonomy was assigned to ASVs using classify-sklearn (via q2-feature-classifier) against the Greengenes 13_8 99% OTUs reference sequences^31^. This classifier was recently shown to achieve similar precision and recall to the RDP classifier^32^ at the genus level on 15 mock community data sets.^33^

### Statistics

Differences in baseline characteristics between the subsample and the parent trial, or between treatment arms, were tested using chi-square tests for categorical variables and t-tests or Wilcoxon rank-sum tests for continuous variables. The difference between the freezer storage time in each treatment arm was tested using a linear mixed effects model, to account for the correlation induced by the baseline and end-of-study samples from the same subject. The association between freezer storage time and microbiome composition was tested using a Spearman correlation coefficient for baseline and end-of-study samples separately. To test for differences in microbiome composition, we performed Principle Coordinate Analysis (PCoA) based on four distance metrics (weighted UniFrac, unweighted UniFrac, Bray-Curtis, and Jaccard). Components of variance was used to estimate the between-patient versus within-patient intraclass correlation coefficient for each microbiome measure. We then computed the change (in direction and magnitude) in the first principal coordinate axis (PC1) for each subject between their pre-treatment and post-treatment samples. The average change in PC1 for each treatment group, overall and stratified by sex, was tested for difference from zero using a one-sample t-test with Benjamini-Hochberg false discovery rate (FDR) correction.^34^ We additionally applied pairwise tests to determine if UDCA treatment was associated with changes in gut microbial community richness (i.e., changes in the number of bacterial taxa present in the community). This was performed by comparing change in Observed OTUs and Faith’s Phylogenetic Diversity on a per subject basis in the two treatment groups.

We performed ANCOM^35^ and Wilcoxon signed-rank tests comparing species abundance at baseline and end-of-study in both UDCA-treated and placebo groups. ANCOM tests were performed to assess differences within the whole bacterial community in each arm separately. Wilcoxon signed-rank tests were additionally performed on 18 individual bacterial genera, the order *Bifidobacteriales*, and the ratio of the *Firmicutes* to *Bacteroidetes* phyla abundances, all of which have been previously associated with CRC or its risk factors.

Associations between change in each microbiome measure and adenoma outcome (any adenoma or advanced adenoma) were tested in each arm separately using Poisson regression, adjusted for sex, age, aspirin use, baseline microbiome measure, and an indicator for whether a participant’s paired baseline and end-of-study DNA samples were processed in different batches. Potential interactions between microbiome measures and UDCA on recurrence were tested using likelihood ratio tests. These statistical tests were performed with Stata 14.2 (StataCorp, College Station, TX).

## RESULTS

### Participant characteristics

Characteristics of the 401-participant subgroup with complete sequence data and adenoma outcome status were compared to participants in the parent trial not included in the microbiome study, by treatment assignment (Table 1). The placebo subsample had fewer aspirin users (chi-square test, *P* = 0.004), the largest adenomas (Wilcoxon rank-sum test, *P* = 0.040), and greater adenoma number at baseline (Wilcoxon rank-sum test, *P* = 0.004) than the placebo parent study. Compared to the parent trial, the UDCA subsample included more male participants (chi-square test, *P* = 0.016). Within the subsample, the UDCA arm included more males (chi-square test, *P* = 0.024) and more aspirin users (chi-square test, *P* = 0.003) than the placebo arm.

**Table 1.** Baseline characteristics of participants in the subsample compared to the parent trial, by treatment arm.

| Variable | Placebo arm |  | UDCA arm |  |
| --- | --- | --- | --- | --- |
|  | Subsample<br>(n = 203) | Parent trial<br>(n = 421) | Subsample<br>(n = 198) | Parent trial<br>(n = 463) |
| Age, mean ± SD | 66.5 ± 8.0 | 66.3 ± 8.5 | 66.2 ± 8.9 | 66.0 ± 8.6 |
| Male, n (%) | 133 (65.5) | 280 (66.5) | 150 (75.8) | 307 (66.3) |
| White, n (%) | 188 (94.0) | 388 (93.7) | 189 (96.9) | 426 (94.3) |
| Education (y), mean ± SD | 13.9 ± 2.3 | 14.1 ± 2.3 | 14.1 ± 2.3 | 13.9 ± 2.2 |
| Ever smoker, n (%) | 134 (69.1) | 293 (71.6) | 125 (64.1) | 314 (69.8) |
| Current smoker, n (%) | 21 (10.3) | 57 (13.5) | 23 (11.6) | 56 (12.1) |
| BMI (kg/m<sup>2</sup>), mean ± SD | 28.4 ± 4.7 | 28.1 ± 4.8 | 28.0 ± 4.9 | 28.1 ± 4.8 |
| Aspirin use, n (%) | 39 (19.2) | 127 (30.2) | 64 (32.3) | 124 (26.8) |
| Family history of CRC, n (%) | 66 (32.5) | 115 (27.3) | 57 (28.8) | 111 (24.0) |
| Previous polyp, n (%) | 77 (40.1) | 189 (48.6) | 94 (48.7) | 209 (48.1) |
| Largest adenoma (mm), mean ± SD; median | 9.6 ± 6.3; 8 | 8.4 ± 5.4; 7.5 | 8.9 ± 5.4; 8 | 8.7 ± 5.4; 8 |
| Number of adenomas, mean ± SD; median | 1.6 ± 0.9; 1 | 1.5 ± 0.8; 1 | 1.7 ± 1.1; 1 | 1.6 ± 0.9; 1 |
| Proximal adenomas, n (%) | 113 (55.7) | 227 (54.2) | 112 (56.6) | 260 (56.3) |
| Villous component to adenoma, n (%) | 46 (22.7) | 78 (18.5) | 33 (16.7) | 106 (23.0) |
| High-grade dysplasia, n (%) | 21 (10.3) | 35 (8.3) | 19 (9.6) | 38 (8.2) |

Missing data: race, n = 24 (1.9%); education, n = 29 (2.3%); ever smoker, n = 37 (2.9%); BMI, n = 29 (2.3%); previous polyp, n = 76 (5.9%); largest adenoma, n = 1 (0.1%); proximal adenoma, n = 3 (0.2%); villous histology, n = 2 (0.2%)

### Microbiome composition is not correlated with storage time

After separation from fecal water, solid stool samples used in this study were stored at −80°C for varying lengths of time before microbiome sequencing. Baseline samples were stored for an average of 17.2 ± 1.1 years, and end-of-study samples were stored for an average of 14.6 ± 1.1 years. There was no significant difference in storage time by treatment arm (P= 0.22). Furthermore, no significant correlations were observed between storage time and any of the diversity metrics at baseline or end-of-study. Lack of evidence for storage time effects on these measures is in agreement with published studies supporting long-term freezing as an effective preservation method for studies of microbiome composition.^36^

### Microbiome changes in response to UDCA treatment

PCoA based on unweighted UniFrac distance between samples does not illustrate a clear difference between baseline and end-of-study microbial communities in either treatment group (Figure 1A). Distances between paired samples from the same subject were smaller than distances between samples from different subjects in both treatment groups (Figure 1B). Intraclass correlation coefficients estimated separately for each of the four beta-diversity metrics ranged from 0.50 to 0.68 for the placebo group, and from 0.39 to 0.73 for the UDCA group (Figure 1B). There was no clear pattern of change in composition between the UDCA and placebo arms in terms of the magnitude of the four measures applied to assess microbial community composition (U = 19292.00, P = 0.244) (Figure 1C), suggesting that both treatment groups experience a similar amount of microbiome change between baseline and end-of-study.

**Figure 1:**
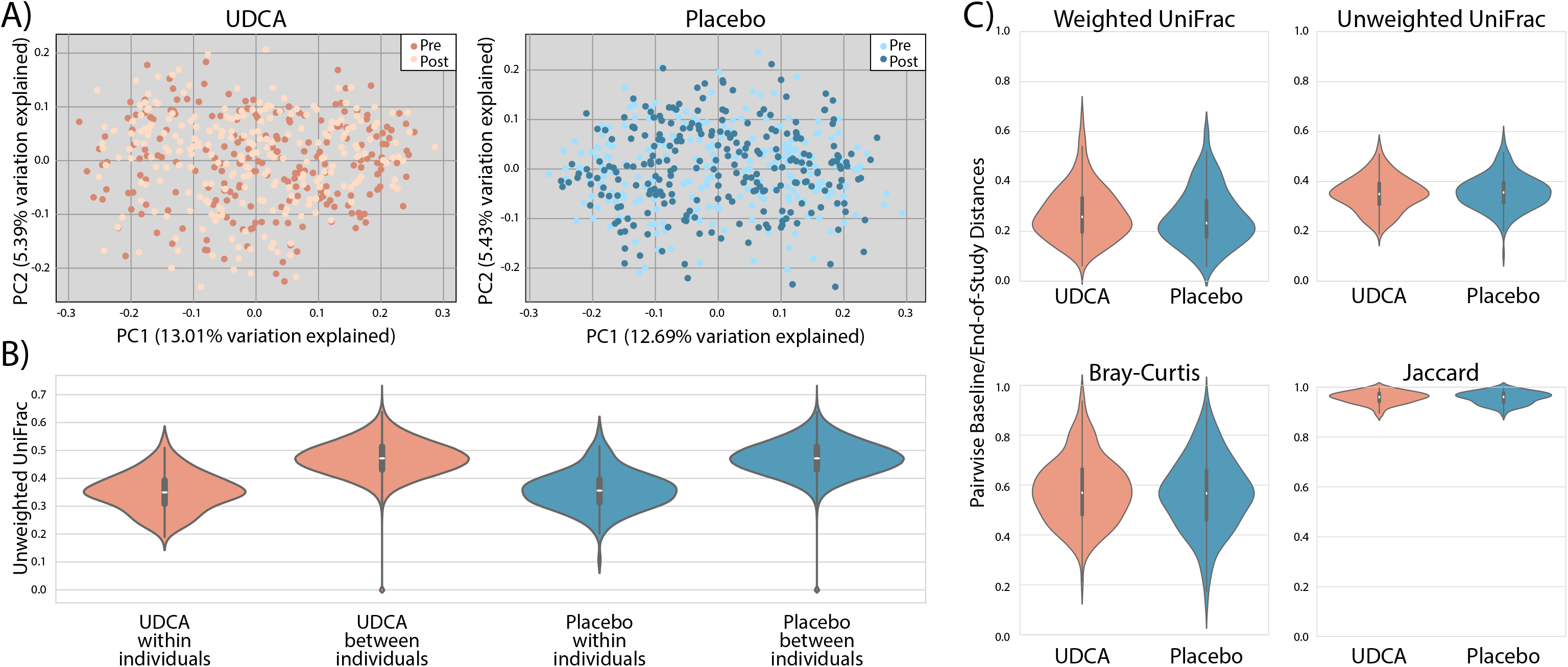
A) PCoA plots for UDCA and placebo groups with pre and post samples (light and dark, respectively). B) Violin plots illustrate the full distribution of data for different values of unweighted UniFrac distances within and between individuals. Marker for the median (center point), interquartile range (box), and 1.5 interquartile range (whiskers) are included. Distances within individuals are significantly less than distances between individuals. C) Violin plots depict the magnitude of change in microbiome composition between baseline and end-of-study in UDCA and placebo groups. The magnitude of change did not differ significantly between the treatment groups for any of these metrics.

Given the amount of microbiome changes from baseline to end-of-study appeared similar between placebo and intervention, we next tested whether individuals in either arm experienced changes in their microbiome that were more similar to one another. Paired one-sample t-tests were used to identify consistent changes across individuals in four microbial community distance metrics (Figure 2A-D) and two microbial community richness metrics (Figure 2E-F). In this analysis, UDCA treatment was associated with a shift in microbial community distance metrics according to PC1 of unweighted UniFrac (t = −4.393, *P* < 0.001) distance, and PC1 of both unweighted (Jaccard: t = −5.697, *P* < 0.001) and weighted (Bray-Curtis; t = −2.699, *P* = 0.035) non-phylogenetic metrics. These shifts were not observed in the placebo arm. These results suggest that while gut microbial communities changed by a similar degree in both UDCA and placebo groups (Figure 1C), individuals in the UDCA arm experienced changes that were more similar to each other (i.e. ‘UDCA-associated’) than those in the placebo arm (Figure 2B-D). For gut microbial community richness (i.e., changes in the number of bacterial taxa present in the community), Observed OTUs and Faith’s Phylogenetic Diversity were computed on a per-subject basis in each arm (Figure 2 E-F). The average change was not significantly different from zero in either arm for either measure (all *P* > 0.05). Therefore, despite UDCA-induced changes in overall community composition, we found no evidence that UDCA treatment significantly altered gut microbial community richness. In other words, the significant compositional changes observed with UDCA treatment support alterations to relative abundance and even presence/absence of microbial species, but not the number of different types of organisms present in the gut microbiome.

**Figure 2:**
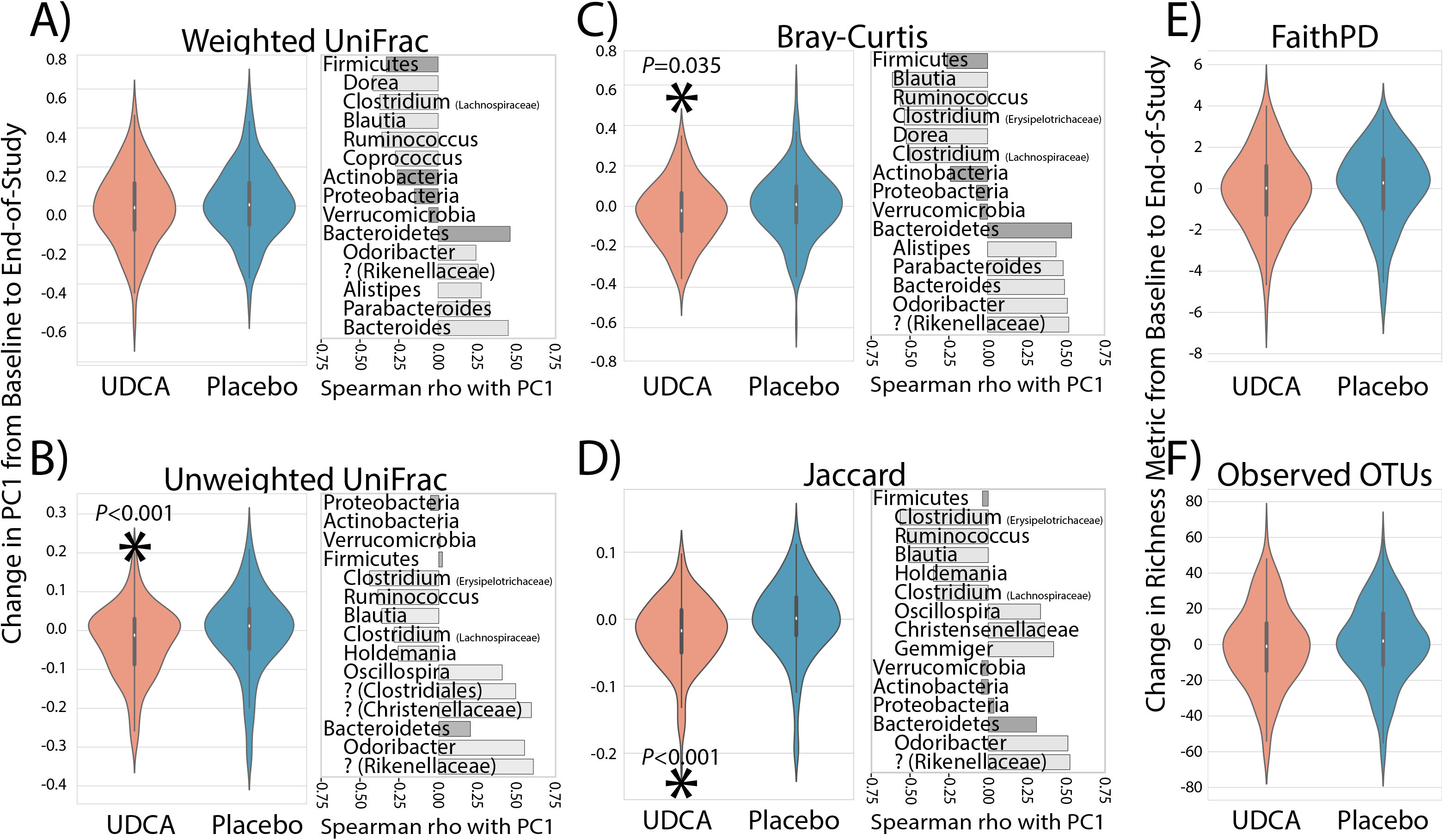
Pairwise changes in PC1 between baseline and end-of-study samples (left panels) and correlation with taxonomic changes (right panels) shown for phyla (dark gray bars) and genera (light gray bars). Question marks indicate unknown genera and include the most specific known taxonomic association in parentheses. A-D) Change in PC1 for microbial community distance metrics in each treatment arm. E-F) Change in microbial community richness metrics in each treatment arm. Statistically significant comparisons are indicated with an asterisk and p-value.

Because UDCA treatment was shown to be protective against the development of adenoma in males but not females in the parent trial,^17^ we next explored results stratified by sex (Figure 3A-F). Using a pairwise approach, two of the six microbiome measures showed a statistically significant change with UDCA treatment in males [unweighted UniFrac (t = −4.393, *P* < 0.001) and Jaccard (t= −5.234, *P* < 0.001)]. For females, none of the metrics showed a significant change with UDCA, likely due to the smaller sample size (48 women versus 150 men), as the mean change in PC1 was the same for females and males. As in the total sample, no systematic changes were observed for males or females in the placebo arm.

**Figure 3:**
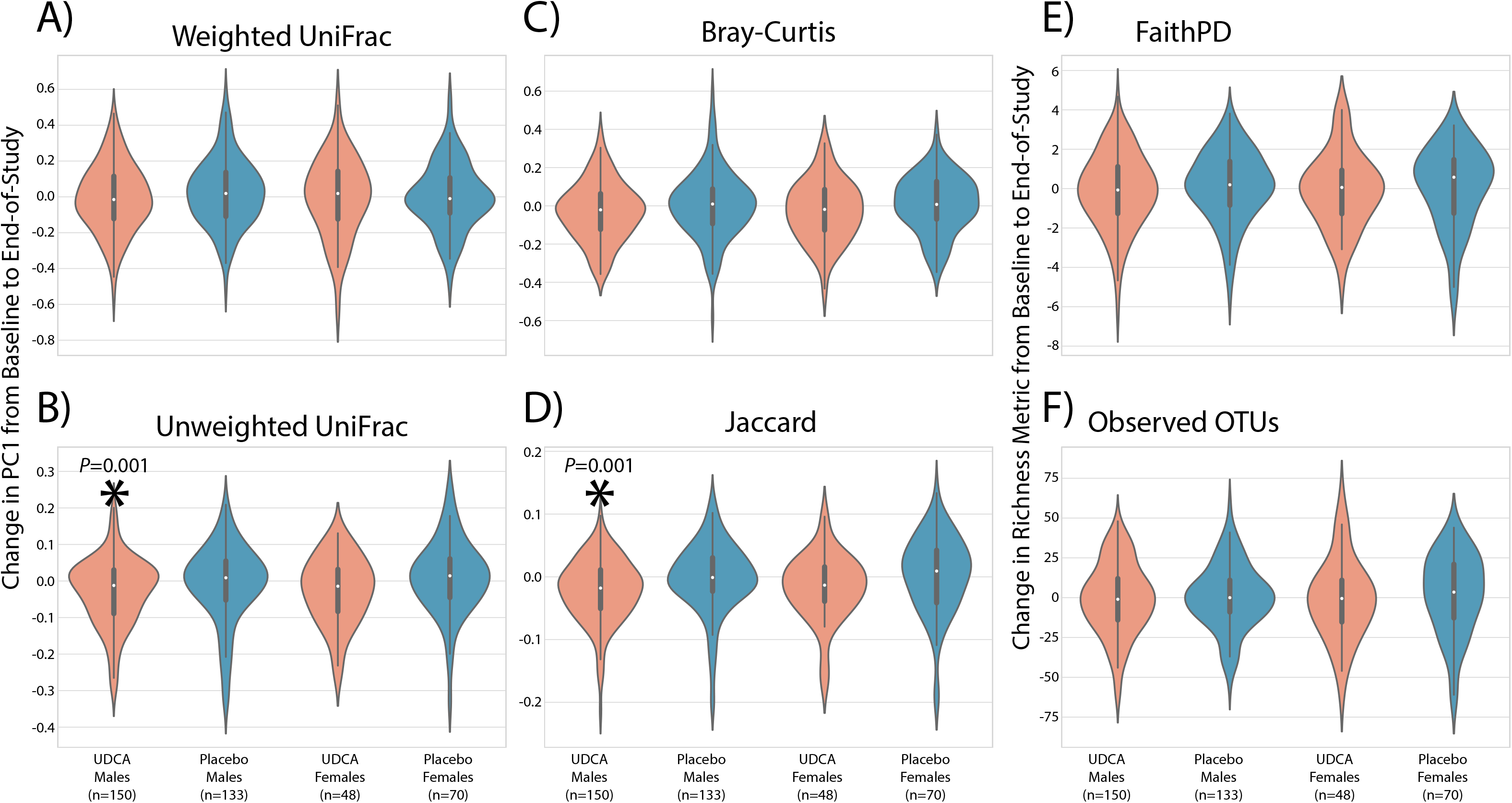
Pairwise changes between baseline and end-of-study samples stratified by treatment arm and sex. A-D) Change in PC1 for microbial community distance metrics. E-F) Change in microbial community richness metrics. Statistically significant comparisons between treatment arms are indicated with an asterisk and p-values.

With the observed changes in community composition in response to UDCA treatment, we were interested in identifying bacterial taxa that exhibited abundance changes. ANCOM tests indicated that no bacterial genera or ASVs consistently differed between baseline and end-of-study measurements in the placebo group. In the UDCA treatment arm, ANCOM tests on all ASVs showed that the relative abundance of *Faecalibacterium* decreased between baseline and end-of-study. Paired Wilcoxon signed-rank tests were additionally performed on 18 individual bacterial taxa that contain species or strains previously associated with CRC,^37^ as well as for the genus *Bifidobacterium* (see **Supplemental Table 1**). Of these, *Streptococcus* (FDR-corrected P = 0.003), *Escherichia* (FDR-corrected P = 0.003), and *Bilophila* (FDR-corrected P = 0.012) were found to have increased significantly, while *Fusobacterium* (FDR-corrected P = 0.049) decreased in relative abundance between baseline and end-of-study in UDCA-treated subjects. There were no significant changes for these genera in the placebo arm (all FDR-corrected P > 0.05). We additionally tested whether the ratio of the Firmicutes to Bacteriodetes phylum abundances changed with treatment using Wilcoxon signed-rank tests, but did not find evidence for this in either treatment group (UDCA: W=10369.5, FDR-corrected P = 0.57; placebo: W=9081.0, FDR-corrected P = 0.13).

### Change in microbiome and adenoma recurrence

We next assessed whether UDCA-associated changes in community composition, when controlled for the baseline value, were associated with adenoma development. We found no evidence that change in any of the four microbial community distance metrics from baseline to end-of-study were associated with risk of adenoma in either treatment arm (all P> 0.05) even after considering effects by sex separately. For the specific ASVs that were shown to increase with UDCA treatment (i.e., *Streptococcus, Escherichia, Bilophila* and *Fusobacterium)*, we found no evidence of association between any of these ASVs and adenoma outcome in either the placebo or UDCA arm (all P >0.05).

## DISCUSSION

Utilizing six metrics of microbiome diversity and richness, we assessed whether daily oral UDCA (6-8 kg/m^2^) given for an average of 3 years for the prevention of colorectal adenoma significantly changed the gut microbiome. Secondarily, we investigated whether change in these microbiome measures were associated with adenoma risk by treatment arm considering sex as a modifying factor of UDCA chemoprevention benefit. Our results show participants randomized to UDCA exhibited non-random changes in their microbiome diversity. However, the UDCA associated changes were not associated with any chemopreventive action of UDCA for adenoma risk. We observed no significant effect of UDCA on species richness (number of observed ASVs, a corollary of the number of species present) or phylogenetic richness of microbial communities nor UDCA-related changes in abundance-weighted UniFrac phylogenetic diversity metrics (which is biased toward detecting changes in distantly related community members that are present in high relative abundance, discussed below). Further, while the observed effects on the microbiome reached significance in the larger sample size of men, the overall pattern of change was similar for women. Our results do not support UDCA effects on the microbiome as a mediator of chemopreventive activity nor do we find evidence of differential effects of UDCA on the gut microbiome by sex as an explanation for our previous finding of chemopeventive effects of UDCA for adenoma in men but not women.

Microbial communities in participants randomized to UDCA differed in their composition, particularly in PC1, as measured by unweighted but not weighted UniFrac. Unweighted UniFrac is a measure of the degree of phylogenetic similarity between two microbial communities not considering abundance of ASVs, and, hence, is equally sensitive to differences in low- and high-abundance ASVs. In contrast, weighted UniFrac accounts for ASV abundances when comparing microbial community composition between samples and is therefore more sensitive to detecting changes in high-abundance ASVs. Both UniFrac metrics are designed to up-weight changes in phylogenetically dissimilar ASVs relative to phylogenetically similar ASVs. Thus, our observation of a significant change in unweighted UniFrac and no significant change in weighted UniFrac suggests that overall distantly related, but lower abundance, ASVs changed with UDCA.

Bray-Curtis (abundance-weighted) and Jaccard dissimilarity measures are the non-phylogenetic analogs to weighted and unweighted UniFrac, respectively; they measure the degree to which two microbial communities share ASVs, rather than the degree of phylogenetic relatedness between communities. We observed a significant change in Bray-Curtis distance. Together with no significant change in weighted UniFrac, our results suggest that high abundant, phylogenetically similar ASVs are changing with UDCA. As Jaccard distance is not phylogenetically or abundance weighted, it limits interpretation of the results and implies only that some ASVs are changing. By comparing our results across these four metrics, we can gain some insight into categories of microbial community members that are changing in response to UDCA. Specifically, our results suggest that distantly related, low-abundance ASVs as well as closely related, high-abundance ASVs (but not distantly related, high-abundance ASVs) are changed with UDCA treatment.

To gain insight on how the observed UDCA changes related to changes in the taxonomic composition of the gut microbiome, we evaluated the bacterial phyla and genera most strongly associated with PC1. This was achieved by computing Spearman correlation coefficients between the phyla and genera that were observed at least one time in at least 50% of the 802 pre- and post-treatment microbiome samples. For most metrics, change in PC1 was associated with an increase in the Bacteriodetes relative abundance and a decrease in the Firmicutes relative abundance with UDCA treatment. The Bacteriodetes and Firmicutes are two dominant microbial phyla comprising the gut microbiome. These common bacteria in the human gut and their ratio to each other have been suggested to reflect dietary pattern and overall balance of the gut microbiome. For example, a high Firmicutes to Bacteroidetes ratio has been associated with consumption of the Western diet^17^ and with adverse metabolic changes that occur with obesity.^38, 39^ In contrast, a low Firmicutes to Bacteroidetes ratio has been associated with reduced gut biodiversity^40^ and observed in patients with inflammatory bowel disease.^41^ While the relative abundance of all Firmicutes and Bacteroidetes did not change in either treatment group, individual Firmicutes taxa tended to decrease while Bacteroidetes taxa increased and were associated with changes along the PC1 axis. As such, the decreases in Firmicutes and increases in Bacteroidetes with UDCA may reflect positive effects of UDCA on the gut microbiome.

UDCA-associated increases in species of *Streptococcus, Escherichia*, and *Bilophila* and decreases in *Fusobacterium* are notable in context of reported associations between different members of these genera and CRC. An increase in Bilophila is biologically consistent with earlier studies, including our own, showing that UDCA led to increases in the levels of DCA in aqueous and solid stool fractions, with evidence that UDCA may enhance fecal bile acid levels through inhibitory effects on 7-α-dehydroxylation of cholic acid. As such, expansion of *Bilophila* would be expected but perhaps not desirable given pro-inflammatory effects of *Bilophila wadsworthia* in mice. Increases in members of the genera *Streptococcus* and *Escherichia* with UDCA may similarly reflect response to changes in the bile acid pool in stools of UDCA subjects. At the 16S RNA level we are unable to assess effects on select strains of bacteria. For example, we are unable to determine the effects of UDCA on streptococcal lactic acid bacteria thought to have anti-mutagenic/anti-cancer properties in human intestine,^42^ from subspecies of *Streptococcus gallolyticus* that have been associated with colon cancer proliferation and growth.^43^ Importantly, we are unable to test for any UDCA effect on *E. coli* strains harboring the polyketide synthase *(pks)* genomic island, which encodes for the genotoxin colibactin, and has been identified in cancer and inflammatory bowel disease and shown to promote tumor development in inflammatory mouse models.^44, 45^ Interesting is the observed UDCA reduction in *Fusobacterium spp*. Several studies have suggested a link between *Fusobacterium spp*. and CRC with interest in *F. nucleatum*. Most recently, this association has been suggested to reflect a ‘passenger’ role where *F. nucleatum* expands in numbers in response to an environment that favors CRC as opposed to a direct causal role^46^ explaining the failure of *F. nucleatum* strains identified in patients to promote colonic tumors in mouse models. Whether UDCA-associated decreases in *Fusobacterium spp*. include a change in *F. nucleatum* warrants more-selective sequence analysis.

Longitudinal variation of the gut microbial community within individuals is expected^47^ and the degree of variation fluctuates between individuals.^48, 49^ This variation, along with the high intraclass correlation coefficients observed in our study and evidence that components of the microbiome are highly individualized,^50^ are significant limiting factors for the detectability of modest effects of medical treatment on the microbiome in all but extreme cases such as vancomycin treatment or fecal microbiota transplant. As such, despite being one of the largest studies of drug effects on the microbiome in the randomized setting, we are unable to rule out modest effects of UDCA on the microbiome as a mechanism of drug effect on colorectal adenoma development.

---

Supplemental Table 1. Wilcoxon signed-rank test comparison comparing relative abundance of carcinogenesis-associated taxa pre- and post-treatment in UDCA-treated subjects

## Abbreviations

UDCA: Ursodeoxycholic acid
CRC: Colorectal cancer
DSS: Dextran sodium sulfate
AOM: Azoxymethane
PBC: Primary biliary cirrhosis
OTU: Operational taxonomic unit
ASV: amplicon sequence variant

